# How well do routine molecular diagnostics detect rifampicin heteroresistance in *Mycobacterium tuberculosis*?

**DOI:** 10.1101/632851

**Authors:** Kamela C. S. Ng, Philip Supply, Frank G. J. Cobelens, Cyril Gaudin, Julian Gonzalez-Martin, Bouke C. de Jong, Leen Rigouts

## Abstract

Rifampicin heteroresistance – where rifampicin-resistant and -susceptible tuberculosis (TB) bacilli co-exist – may result in failed standard TB treatment and potential spread of rifampicin-resistant strains. Detection of rifampicin heteroresistance in routine rapid diagnostic tests (RDTs) allows for patients to receive prompt and effective multidrug-resistant-TB treatment, and may improve rifampicin-resistant TB control.

The limit of detection (LOD) of rifampicin heteroresistance for phenotypic drug susceptibility testing by the proportion method is 1%, yet is insufficiently documented for RDTs. We therefore aimed to determine, for the four RDTs (XpertMTB/RIF, XpertMTB/RIF Ultra, GenoTypeMTBDR*plus*v2.0, and GenoscholarNTM+MDRTBII), the LOD per probe and mutation, validated by colony-forming-unit-counting and targeted deep sequencing (Deeplex-MycTB).

We selected one rifampicin-susceptible and four rifampicin-resistant strains, with mutation D435V, H445D, H445Y, and S450L respectively, mixed them in various proportions in triplicate, tested them with each RDT, and determined the LODs per mutation type. Deeplex-MycTB revealed concordant proportions of the minority resistant variants in the mixtures. The Deeplex-MycTB-validated-LODs ranged from 20-80% for XpertMTB/RIF, 20-70% for Xpert Ultra, 5-10% for GenoTypeMTBDR*plus*v2.0, and 1-10% for GenoscholarNTM+MTBII for the different mutations.

Deeplex-MycTB, GenoTypeMTBDR*plus*v2.0, and GenoscholarNTM+MDRTBII, provide explicit information on rifampicin heteroresistance for the most frequently detected mutations. Classic Xpert and Ultra report rifampicin heteroresistance as rifampicin resistance, while Ultra may denote rifampicin heteroresistance through ‘mixed patterns’ of wild-type and mutant melt probe melt peak temperatures.

Overall, our findings inform end-users that the threshold for reporting resistance in case of rifampicin heteroresistance is the highest for Classic Xpert and Ultra, to resolve phenotypic and genotypic discordant rifampicin-resistant TB results.

## INTRODUCTION

Resistance to rifampicin (RIF) – the most potent core drug in the standard tuberculosis (TB) treatment regimen (1) – is a major barrier to TB control. In 2017, 71% of global RIF resistant-TB (RR-TB) cases were not diagnosed (2). The diagnosis of RR-TB may be complicated by RIF heteroresistance, observed in patient samples where RR and RIF-susceptible (RS) strains co-exist (3, 4), which may be missed and diagnosed as RS because of detection limits. RIF heteroresistance may arise from an existing resistant clonal subpopulation or from a mixed infection of independent strains with RR and RS profiles. Such heteroresistance, also known as ‘unfixed’ resistance, precedes full blown resistance (‘fixed’ resistance, 100% RR) as a result of further resistance selection under treatment (4–7). Failure to detect these minority resistant variants can thus result in unsuccessful treatment and potential spread of RR-TB strains (3, 4, 8).

The World Health Organization currently endorses use of rapid diagnostic tests (RDTs) for timely detection of RR-TB strains: XpertMTB/RIF (classic Xpert), Xpert Ultra (Ultra), and the Line Probe Assays (LPAs) GenoTypeMTBDR*plus*v2.0 (LPA-Hain) and Genoscholar NTM+MDRTBII (LPA-Nipro) (9–11). Among these RDTs, only the LPAs are currently known to explicitly detect RIF heteroresistance in case of mixtures with mutations covered by the assay, exemplified by both wild-type and mutant bands being present, also known as ‘mixed patterns’ (4, 12).

In this study, we define RIF heteroresistance limit of detection (LOD) as the minimum proportion of mutant bacilli in a total mycobacterial population present in a sample, needed for RIF resistance to be detected (4). It is known that phenotypic drug susceptibility testing by the proportion method determines at least 1% resistant subpopulation in clinical samples (4, 13, 14). In the abovementioned RDTs however, the RIF heteroresistance LOD in association with the specific *rpoB* mutation, is insufficiently documented. In the case of classic Xpert, previous studies report LOD values ranging from 65 to 100% (15, 16), for Ultra, the first validation study conducted by the manufacturer presented LODs only for mutations L430P, H445N (20-40%), and S450L (5-10%) (17), whereas LODs of Genoscholar NTM+MDRTB II have not been reported yet.

High coverage depths achieved through pre-selected amplified genes allow targeted deep sequencing to capture and quantify minority resistant variants of *Mycobacterium tuberculosis* mutants and detect RIF heteroresistance with high sensitivity (18, 19). As an example of such an approach, Deeplex®-MycTB (Genoscreen, France; Deeplex) employs ultra-deep sequencing of a single, 24-plexed amplicon mix to detect drug resistance-associated mutations in *M. tuberculosis* complex (MTBC) strains, in addition to mycobacterial species identification and MTBC strain genotyping, with a 24-48 hours turnaround time starting from smear positive clinical samples or primary cultures. Among the 18 main gene targets associated with 1^st^ and 2^nd^– line drug resistance included in Deeplex-MycTB, the *rpoB* gene – associated with RR - is covered by two amplicons of which one comprises the main mutation hotspot region also known as the rifampicin resistance determining region (RRDR) (20).

Precise documentation of LODs for most relevant *rpoB* mutations and for the state-of-the-art RDTs is necessary for timely and more accurate identification of RIF heteroresistance and prompt initiation of appropriate treatment. Therefore, we determined the LODs of classic Xpert, Ultra, LPA-Hain, and LPA-Nipro for detecting RIF heteroresistance linked with RR mutations S450L, D435V, H445D, and H445Y, in relation with the different probes used in each RDT. These four mutations are most frequently detected in association with RR-TB in the global MTBC strain population according to large-scale studies (10, 21). We used simulated mixtures of selected, cultured RR and RS-TB strains, at ratios initially based on colony-forming-unit (CFU) counts after McFarland standardization of the bacillary suspensions. Targeted deep sequencing by Deeplex-MycTB was used as reference for quantitative assessment of the RR:RS ratios.

## MATERIALS AND METHODS

### Selection of strains

We selected one RS- and four RR-*M. tuberculosis* strains from the Belgian Coordinated Collection of Microorganisms hosted in the Institute of Tropical Medicine Antwerp (22), on the basis of the presence of mutations confirmed by Sanger sequencing and captured by mutation probes of LPA-Hain and LPA-Nipro: the Beijing 2.2.1.1 strains TB-TDR-0090 (ITM-041208 with *rpoB* mutation S450L (S531L in *E.coli* numbering)) and TB-TDR-0100 (ITM-041220, D435V (D516V)); the LAM 4.3.4.2 strains TB-TDR-0036 (ITM-000930, H445D (H526D)) and TB-TDR-0131 (ITM-041289, H445Y (H526Y)) (23). The RS strain was the Euro-American lineage 4.9 TDR-0140 (ITM-091634, *rpoB* wildtype).

### Bacillary suspensions and baseline colony-forming-unit (CFU) counting

We prepared two batches of McFarland standard 1 (24, 25) suspensions for each RR and the RS strain.. To check if numbers of bacilli were similar among the cultures after McFarland standardization, we performed CFU counting by spread plating of serial dilutions until 10^−4^-10^−6^ (Figure S1 a). From each dilution, 100µl was plated in triplicate on Dubos agar plates that were sealed with a double layer of parafilm, placed in ziplock bags and incubated at 37°C for four weeks, before colony counting. The first batch was used to prepare RR:RS mixtures for testing by classic Xpert, LPA-Hain, and LPA-Nipro, and the second batch was prepared for assessment by Ultra, which was only released after initial testing.

### Simulation of RIF heteroresistance

RIF heteroresistance was simulated for each mutation type, by mixing McFarland standard 1 suspensions of the RS and respective RR strains in triplicate in the following proportions (R:S): 0:100, 1:99, 5:95, 10:90, 20:80, 30:70, 40:60, 50:50, 60:40, 70:30, 80:20, and 100:0 (Figure S1 b), and vortexing the mixtures for 20 seconds. Replicate 3 of each RR:RS mixture per batch was tested by targeted deep sequencing (Deeplex-MycTB), the results of which served for cross-validation of variant quantification.

### Subjection of RR:RS mixtures to RDTs

We subjected mixtures of RR- and RS-TB bacillary suspensions to classic Xpert and Ultra (10, 11), and thermolysates to LPA-Hain and LPA-Nipro following manufacturer’s instructions. We recorded the LODs and corresponding RDT probe reaction per mutation type, in comparison with values cross-validated by CFU counts and variant quantification with Deeplex-MycTB. First reading of the LPA strips was done by the person who prepared the mixtures and performed the tests, while second reading was done by a colleague who was blinded to the sample information to ensure objective reading of raw results. Additionally, for LPA-Nipro, the Genoscholar Reader - a mobile application developed by Nipro (Osaka, Japan, https://itunes.apple.com/by/app/genoscholar-reader/id1149733183?mt=8) was utilized.

In their standard reporting, Classic Xpert and Ultra (Supplemental File Figure S4 a) report RIF heteroresistance, above or equal its LOD, as RR. Users can indirectly infer RIF *hetero*resistance from Ultra data, as shown by the simultaneous presence of wild-type and mutant (Mut) melt peak temperatures at the ‘Melt Peaks’ tab (Supplemental File Figure S4 b). When generating the results in portable document format, users must tick the ‘Melt Peaks’ box to include the melt peak temperatures associated with each wild-type and Mut melt probe, which may denote RIF heteroresistance, in the extended report (Supplemental File Figure S4 b). Full resistance is detected by presence of both rpoB4A and rpoB3 Mut melt probes for mutation S450L, whereas RIF heteroresistance is detected only by the rpoB4A Mut melt probe in combination with the corresponding wild-type melt probe (Figure 1).

**Figure 1.**
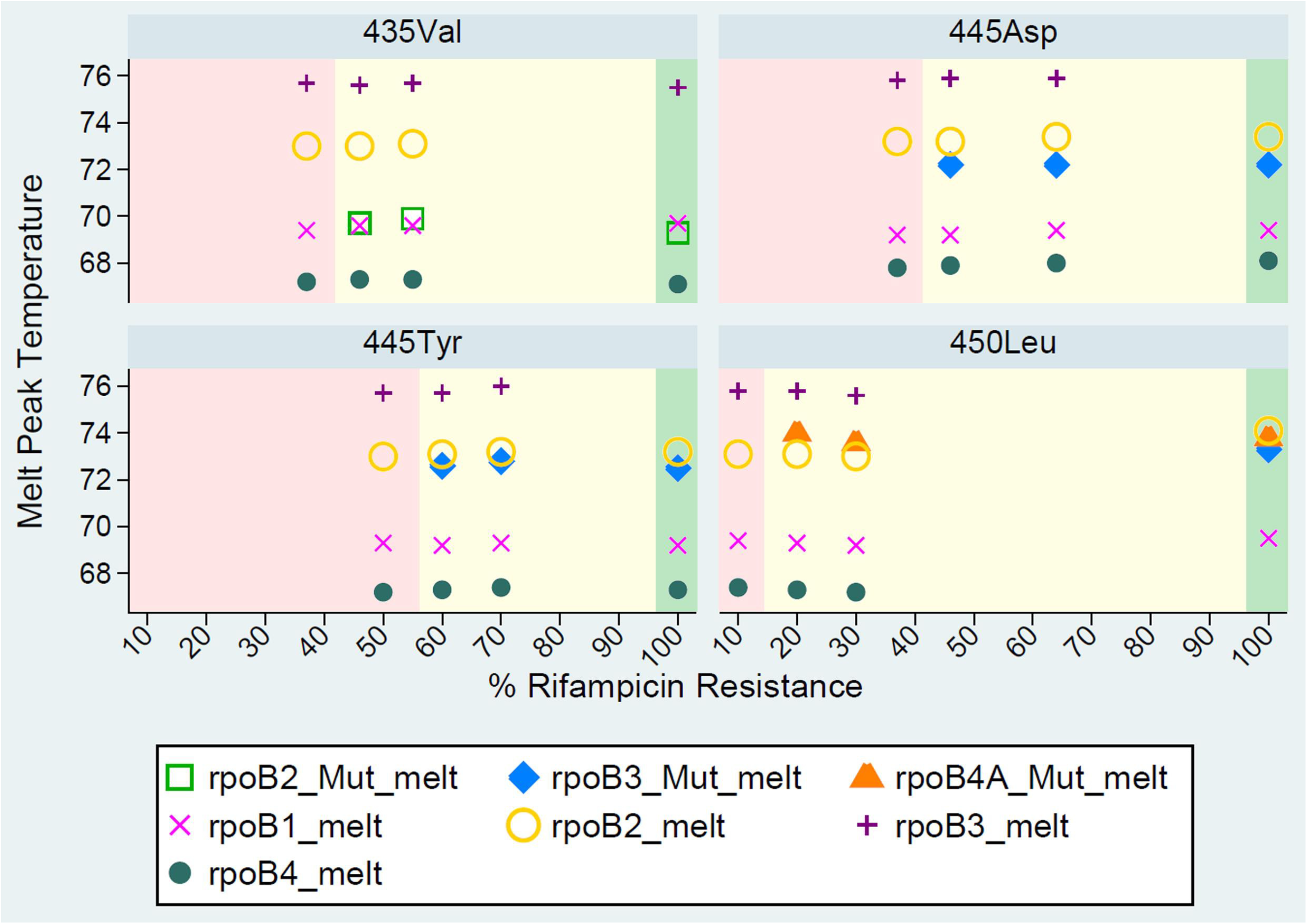
Presence/absence of Xpert Ultra wild-type and mutant melt probes detected for the rifampicin-resistant(RR):rifampicin-susceptible(RS) mixtures per mutation type at various proportions of the minority resistant variants: at the limit of detection (LOD) reported as (hetero)resistant (shaded in yellow), below LOD reported as RS (shaded in pink), and at full RR reported as resistant (shaded in green).

### Quantitative assessment of mutant proportions by Deeplex-MycTB

Per batch, we quantitatively assessed proportions of resistant subpopulations in replicate 3 of the prepared RR:RS mixtures by Deeplex-MycTB. These proportions were determined by calculating the mean percentages of minority resistant variants across all mutation positions borne by each of the RR strains in the *rpoB* gene and other gene targets depending on the strain genetic background.

Thermolysates of the RR:RS mixtures prepared as previously described (26) were subjected to amplicon deep sequencing, using Deeplex-MycTB Kits for the amplification according to the manufacturer’s instructions. Replicates 3 of all first batch mixtures were tested with the classical 18-gene target version, while replicate 3 of the second batch with S450L-WT mixtures was tested with a customized version, including 5 gene targets (*rpoB, katG, inhA, fabG1, gyrA*). Amplicons were purified using Agencourt® AMPure® XP magnetic beads (Beckman Coulter, USA) and quantified by fluorescence quantification in 96-well plates on Victor. Paired-end libraries of 150-bp read length were prepared using the Nextera XT DNA Sample Preparation kit (Illumina Inc., San Diego, 160 CA, USA) and sequenced on Illumina MiniSeq using standard procedures. Variant calling was performed using a dedicated, parameterized web application developed by Genoscreen. The nominal threshold for calling minority resistant variants – indicating heteroresistance for drug resistance associated mutations – is set at a minimum of 3% of all reads, after filtering and depending on the coverage depths, to minimize false positive calls due to background technical noise (20, 27, 28). Variants present in lower proportions than 3% in the relevant *rpoB* mutation positions were detected separately from the web application,(29) without application of this nominal threshold in the analysis pipeline.

## RESULTS

After McFarland standardization of the different strain cultures, the mean CFU counts for dilutions 10^−4^-10^−6^ of the three replicates of both first and second batches of the different RS and RR bacillary suspensions were very similar (Table S1 a and b). The proportions obtained from the Deeplex-MycTB analysis were overall consistent with the expected mixture ratios and the relative variation seen among CFU counts, as relative deviations from expected values were limited to 0-16% (Table 1, Figure S2).

**Table 1.** Proportions of minority resistant variants detected among the rifampicin-resistant:rifampicin-susceptible mixtures by targeted deep sequencing (Deeplex-MycTB). The values represent the percentage of sequence reads bearing the indicated *rpoB* mutation. Gray boxes represent proportions below the nominal threshold of 3% for calling minority resistant variants used in the Deeplex-MycTB application.

| RR:RS<br>proportions | Minority resistant variant proportion detected for each specific<br>mutation by Deeplex-MycTB |  |  |  |
| --- | --- | --- | --- | --- |
|  | TB-TDR-0036 | TB-TDR-0090 | TB-TDR-0100 | TB-TDR-0131 |
|  | (H445D/H526D) | (S450L/S531L) | (D435V/D516V) | (H445Y/H526Y) |
| 0:100 | 0.03 | 0.05 | 0.0 | 0.03 |
| 1:99 | 1.3 | 1.3 | 1.6 | 1.0 |
| 5:95 | 7.5 | 5.0 | 6.0 | 4.3 |
| 10:90 | 14.7 | 10.0 | 10.0 | 8.7 |
| 20:80 | 28.6 | 21.7 | 25.2 | 20.8 |
| 30:70 | 39.9 | 28.4 | 35.4 | 26.0 |
| 40:60 | 55.9 | 41.6 | 43.6 | 37.9 |
| 50:50 | 60.0 | 50.1 | 60.6 | 46.0 |
| 60:40 | 69.4 | 61.9 | 67.0 | 57.5 |
| 70:30 | 77.6 | 64.9 | 69.6 | 66.7 |
| 80:20 | 85.5 | 69.8 | 86.2 | 79.0 |
| 100:0 | 98.3 | 99.9 | 98.3 | 98.9 |

We excluded replicate 3 from the first batch of S450L-WT preparation due to substantially high deviation across all mixture ratios, as revealed by Deeplex-MycTB (Supplemental Figure S3). This deviation potentially reflects pipetting variation or bacillary clumping despite similar CFU counts of the respective RR and RS strains. Hence for the S450L-WT mixture, replicate 3 of batch 2 – which was initially only tested by Ultra and Deeplex – was subjected also to classic Xpert and the LPAs, given the good correlation between the two batches and to ensure triplicate testing for all mixtures.

In line with these levels of experimental variation, all *rpoB* variants from 5:95 RR:RS mixtures were called by the Deeplex application in proportions ranging from 4.3% (H445Y) to 7.5% (H445D)(Table 1). When the analysis pipeline was used without applying this threshold, expected variants of 1:99 mixtures were also detected in percentages ranging from 1.0% (H445Y) to 1.6% (D435V), which were above background levels of 0.0 to 0.05% detected in the 0:100 mixtures (i.e. wild type strain only) on these specific sequence positions. The average minimum coverage depth observed from the Deeplex analyses was 1595 reads.

Among the available classical RDTs, LPAs had a lower LOD to detect RIF heteroresistance compared to classic Xpert and Ultra. The proportion of variants required to be detectable through LPA-Hain was 5% for mutation S450L and 5-10% for mutations D435V, H445D, and H445Y (Table S2). LPA-Nipro performed similarly, with 1-5%, 5%, 5-10% and 10% resistant bacilli detected for mutations S450L, H445D, D435V, and H445Y respectively (Table S2). LPA reading results were consistent between the two readers. Additionally, the ‘automated’ Genoscholar reader had similar results with manual reading for mutations H445Y and S450L; whereas it had lower sensitivity for mutations D435V (20% Genoscholar reader vs 5% manual reading) and H445D (10% Genoscholar reader vs 5% manual reading) (Table S3, Figure S5 a-d).

In contrast, Classic Xpert detected mutation S450L only in mixtures with at least 20-40% resistant bacilli, mutation H445D with at least 40-60% and mutations D435V and H445Y with at least 70-80% mutant bacilli (Table S2). Likewise, Ultra required a minimum of 20-30% resistant bacilli to detect mutation S450L, 40-50% for D435V, 40-60% for H445D and 60-70% for H445Y (Table S2; Figure 1).

Notably, in case of S450L at 20% mutant bacilli, only Ultra rpoB4 Mut melt A probe was observed, whereas both rpoB4 Mut melt A and rpoB3 Mut melt probes were present in case of 100% S450L (Figure 1). For mutations D435V, H445D, and H445Y only one Mut melt probe was observed, whether hetero-or fully resistant. The melt peak temperatures did not differ among hetero- or fully resistant populations for all four mutations tested (Figure 1).

## DISCUSSION

Consistent with previous studies, we found that LPA-Hain and LPA-Nipro detect rifampicin heteroresistance better than classic Xpert and Xpert Ultra in samples with sufficient *M. tuberculosis* complex target DNA and subpopulations that carry the most frequently occurring RR-conferring mutations. The consistent CFU counts (Table S1 a and b) and NGS data from targeted deep sequencing (Table 1, Figure S2) did not suggest sub-optimal preparation of the mixtures, supporting the LODs we observed that differed by capturing RDT probe and mutation type. Clearly, Deeplex-MycTB provided useful quantitative information for validating qualitative observations from the RDTs. We also show here that in contrast with classic Xpert, Xpert Ultra may denote rifampicin heteroresistance through ‘mixed patterns’ of wild-type and mutant melt probe melt peak temperatures, which can be leveraged to inform target end-users such as reference laboratory staff and researchers. The different LODs observed can be linked to the inherent detecting mechanisms of the RDTs. Xpert is an automated cartridge-based assay that employs heminested real-time polymerase chain reaction assay and molecular beacon technology in which short overlapping probes bind to the rifampicin resistance-determining region of the wild-type *M. tuberculosis rpoB* gene (15, 30). The line probe assays on the other hand rely on multiplex amplification and reverse hybridization involving both wild-type and mutant probes on a membrane strip (31). The LODs of LPA-Hain were consistent with a previous study that employed version 1 of the LPA-Hain kit for mutations H445Y and S450L (4). The initial visual reading of LPA results was consistent with results of second blinded reading for both LPA-Hain and LPA-Nipro, and Genoscholar Reader for LPA-Nipro, although we observed that the mobile application was less sensitive than visual reading for mutations D435V and H445D.

Classic Xpert performed relatively poorly in detecting heteroresistant mixtures with mutation S450L, which is by far the globally most prevalent allele in RR-TB (10). Nevertheless, the 20-40% LOD we found for classic Xpert was lower (more sensitive) compared to previous studies which recorded 65-90% LOD (15, 16). Ultra performed similarly to classic Xpert in detecting minority resistant variants of mutations S450L and H445D, but did relatively better in capturing those of mutations D435V and H445Y (Table 2). The 20% Ultra LOD for S450L was slightly less sensitive than the 5-10% LOD reported for the same mutation by Chakravorty and colleagues (17) who tested single mixtures of RR and RS DNA. We observed higher (less sensitive) LODs for the non-S450L mutations, consistent with their findings.

**Table 2.** (Range of) limits of detection for rifampicin heteroresistance among triplicates, tested by different rapid diagnostic tests, per mutation type, with quantification of the respective lowest proportion by targeted deep sequencing (Deeplex-MycTB).

| Strain ID | Rifampicin resistance-conferring mutation |  | LPA-Hain | Deeplex | LPA-Nipro | Deeplex | Xpert | Deeplex | Ultra | Deeplex |
| --- | --- | --- | --- | --- | --- | --- | --- | --- | --- | --- |
|  |  |  | %LOD<sup>1,a</sup> | %QP<sup>1,b</sup> | %LOD<sup>1,a</sup> | %QP<sup>1,b</sup> | %LOD<sup>1,a</sup> | %QP<sup>1,b</sup> | %LOD<sup>2,a</sup> | %QP<sup>2,b</sup> |
| TB-TDR-0036 | H445D* | H526D^ | 5-10 | 7.5 | 5 | 7.5 | 40-60 | 55.9 | 40-60 |  |
| TB-TDR-0090 | S450L* | S531L^ | 5 | 5 | 1-5 | 5 | 20-40 | 21.7 | 20-30 | 21.7 |
| TB-TDR-0100 | D435V* | D516V^ | 5-10 | 6 | 5-10 | 6 | 70-80 | 69.6 | 40-50 |  |
| TB-TDR-0131 | H445Y* | H526Y^ | 5-10 | 4.3 | 10 | 8.7 | 70-80 | 66.7 | 60-70 |  |

\*in *M. tuberculosis* numbering; ^in *E.coli* numbering ; Deeplex-MycTB = targeted deep sequencing; RDTs rapid diagnostic tests; LPA-Hain = GenoType MTDRplus v2.0; LPA-Nipro = Genoscholar NTM + MTB II; Xpert = classic Xpert MTB/RIF; Ultra = Xpert MTB/RIF Ultra; <sup>1</sup> = batch 1; <sup>2</sup> = batch 2; <sup>a</sup>Limit of detection (range) among triplicates for the RDTs; <sup>b</sup>quantified proportion of minority resistant variants determined by Deeplex on replicate 3.

Apart from the slight differences in the LODs recorded for classic Xpert and Ultra, we observed that Ultra allows users to infer from raw results on the computer screen – under ‘Melt Peaks’ tab – the phenomenon of heteroresistance (Supplemental File Figure S4 b, Figure 1), which is not possible from classic Xpert data. RIF heteroresistance may be rapidly detected by Ultra through observed melt peak temperatures of both wild-type and Mut melt probes akin to mixed patterns of absent wild-type and developed mutant bands in LPA. Further, in a sample with S450L mutation, observing only the melt peak temperature of rpoB4 Mut melt A probe (vs rpoB4 Mut melt A and rpoB3 Mut melt probes for 100% RR) and corresponding wild-type melt probe may also denote RIF heteroresistance (Figure 1). This information is deemed useful for conducting research on Ultra data and could be practical in the field if export of the melt peak temperatures through the LIS port becomes feasible in future software updates, together with other raw data (e.g. melt peak temperatures of wild-type and Mut melt probes associated with RR mutation). Currently, the raw data of Xpert Ultra are only available directly from the module/computer where it was tested, and cannot yet be automatically extracted and shared through the. gxx files. Further, local staff are not usually trained to interpret melt peak temperatures, as it entails considerable effort for the NTPs to train staff in peripheral settings for such advanced interpretation of raw results. We have formally requested access from the Xpert Ultra manufacturer for automated extraction of melt peak temperatures at central level via non-proprietary connectivity platforms.

Automated capture of the melt peak temperatures by connectivity solutions such as DataToCare (Savics, Belgium), GXAlert (SystemOne, USA), or C360 (Cepheid, USA) will avoid tedious and error prone manual transcription of the values from the computer screen. It may also allow the laboratory staff and in-country expert Xpert Ultra users to interpret the melt peak temperatures and their association with specific RR mutations with support of global expert bodies such as the Foundation for Innovative New Diagnostics and Global Laboratory Initiative so that TB reference laboratories can advise peripheral laboratories and clinics optimally. With possible integration into e-Health patient charts beyond tuberculosis diagnostics, this will allow to transform Xpert Ultra data into usable information for the National TB Programs using a combination of unique patient ID and geographical information (10, 32). This may not only greatly benefit the remote resolution of discordant results to improve patient management, but also aid in building more systematic data on the prevalence and impact of heteroresistance on a programmatic level, critical for improving interventions for patients with confirmed heteroresistance.

Our study has limitations. We performed LPA testing of thermolysates in order to allow ‘optimal’ reading of results as indirect LPA testing increases the intensity of the bands. Routine LPA is commonly done directly on clinical specimens, where background hybridization is more common and can be harder to distinguish from heteroresistance, as both phenomena may produce faint(er) bands which are often disregarded. Thus, the sensitivity for heretoresistance detection that we determined for the LPAs likely represents the upper bound of values achievable in clinical practice. Inclusion of a culture step however, may lose minority subpopulations (33) and bias the ratio of mutant and wildtype populations especially if there is fitness loss, and cause delay in obtaining results. Moreover, the LPAs will not report heteroresistance for mutations not covered by a mutant probe, and are not recommended for testing paucibacillary smear-negative samples due to their lower sensitivity of detecting *M. tuberculosis* compared with Xpert and culture (34–36).

We addressed potential variability resulting from bacillary clumping and pipetting by inclusion of biological replicates, showing differences in LOD of maximum two dilutions amongst replicates with the same mutation.

In conclusion, we report distinct abilities of LPA-Hain, LPA-Nipro, classic Xpert, and Ultra to detect minority resistant variants representing the most common RR-conferring mutations against the quantitative results of Deeplex-MycTB. The LPAs have more sensitive LODs than classic Xpert and Ultra in samples with sufficient *M. tuberculosis* complex target DNA, although they report heteroresistance only for the four most common undisputed mutations – D435V, H445D, H445Y, and S450L. For mutations without confirmatory mutant (MUT) band, such as L452P, RIF heteroresistance cannot be detected, and with faint intensity of the WT band (31), it may be difficult to distinguish between WT and (hetero)resistance. Ultra can detect RR and RIF heteroresistance associated with all *rpoB* mutations within the hotspot, albeit requiring a higher proportion of mutant bacilli than LPA. The LPAs and Deeplex-MycTB provide direct information on the occurrence of RIF heteroresistance, whereas Ultra, after informing that RR was detected, may suggest RIF heteroresistance only through additional examination of wild-type and Mut melt probes and corresponding melt peak temperatures in the raw data on the computer screen or in the generated extended report (Supplemental File Figure S4 b).

The clinical importance of heteroresistance is likely substantial (3), akin to ‘fixed‘, i.e. 100% resistance. The proportion method for phenotypic drug susceptibility testing, which has been around for over half a century, by design, tests for ≥1% resistant subpopulations (4, 13, 14), with strong predictive value for poor treatment outcome, at least for the core drugs, e.g. fluoroquinolones (37) and rifampicin (4). Moreover, samples with ≥5% minority *gyrA* resistant variants were found to have the same minimum inhibitory concentration level as that of samples with 100% fluoroquinolone resistance (L. Rigouts, in press, (37)), while for *rpoB*, the Mycobacterium Growth Indicator Tube phenotypic drug susceptibility testing results were similar for samples with ≥5% minority resistant variants and 100% RIF resistance (4). Taken together, this implicates that in samples with resistant subpopulations of at least 5%, classification of heteroresistance as ‘RR’, even when less sensitive than the proportion method, is probably key, notwithstanding the lack of direct evidence on clinical impact. Our findings can thus inform and guide TB reference laboratory staff, healthcare providers, and researchers, that the threshold for reporting resistance in case of rifampicin heteroresistance is the highest for Classic Xpert and Ultra, to resolve phenotypic and genotypic discordant rifampicin-resistant TB results. Prospective large-scale clinical studies using next generation sequencing approaches (3, 8) are necessary to establish the proportion of mutants that predicts poor outcome of treatment with the specific drug.

## ACKNOWLEDGEMENTS

We thank the Belgian Coordinated Collections of Microorganisms in the Institute of Tropical Medicine Antwerp (http://bccm.belspo.be/about-us/bccm-itm) for the strains tested in this study; and Siemon Gabriëls, Michèle Driesen, Wim Mulders, Jelle Keysers, and Pauline Lempens of the Institute of Tropical Medicine Antwerp Mycobacteriology Unit, and Stéphanie Duthoy and Gaëlle Bisch of Genoscreen for their contribution in the experiments and data analysis.

K.C.S.N. was supported by Erasmus Mundus Joint Doctorate Fellowship grant 2016-1346, and B.d.J. and L.R by an ERC starting grant INTERRUPTB (311725). The funders had no role in the study design, data collection and interpretation, or the decision to submit the work for publication.

Philip Supply was a consultant of Genoscreen.

## Notes

#### Summary of Updates

We have added a figure which may guide end-users on how Xpert Ultra may denote rifampicin heteroresistance through 'mixed patterns' of wild-type and mutant melt probe melt peak temperatures.

## REFERENCES

1. Van Deun A, Decroo T, Piubello A, de Jong BC, Lynen L, Rieder HL. 2018. Principles for constructing a tuberculosis treatment regimen: the role and definition of core and companion drugs. Int J Tuberc Lung Dis 22:239–245.

2. WHO. 2018. Global Tuberculosis Report 2018. Geneva.

3. Ley SD, de Vos M, Van Rie A, Warren RM. 2019. Deciphering Within-Host Microevolution of Mycobacterium tuberculosis through Whole-Genome Sequencing: the Phenotypic Impact and Way Forward. Microbiol Mol Biol Rev 83:e00062–18.

4. Folkvardsen DB, Thomsen VO, Rigouts L, Rasmussen EM, Bang D, Bernaerts G, Werngren J, Toro JC, Hoffner S, Hillemann D, Svensson E. 2013. Rifampin heteroresistance in Mycobacterium tuberculosis cultures as detected by phenotypic and genotypic drug susceptibility test methods. J Clin Microbiol 51:4220–2.

5. Hofmann-Thiel S, van Ingen J, Feldmann K, Turaev L, Uzakova GT, Murmusaeva G, van Soolingen D, Hoffmann H. 2009. Mechanisms of heteroresistance to isoniazid and rifampin of Mycobacterium tuberculosis in Tashkent, Uzbekistan. Eur Respir J 33:368–74.

6. Kimerling ME, Kluge H, Vezhnina N, Iacovazzi T, Demeulenaere T, Portaels F, Matthys F. 1999. Inadequacy of the current WHO re-treatment regimen in a central Siberian prison: treatment failure and MDR-TB. Int J Tuberc Lung Dis 3:451–3.

7. Lienhardt C, Cook SV, Burgos M, Yorke-Edwards V, Rigouts L, Anyo G, Kim SJ, Jindani A, Enarson DA, Nunn AJ, Study CTG. 2011. Efficacy and safety of a 4-drug fixed-dose combination regimen compared with separate drugs for treatment of pulmonary tuberculosis: the Study C randomized controlled trial. JAMA 305:1415–23.

8. Metcalfe JZ, Streicher E, Theron G, Colman RE, Allender C, Lemmer D, Warren R, Engelthaler DM. 2017. Cryptic Microheteroresistance Explains Mycobacterium tuberculosis Phenotypic Resistance. Am J Respir Crit Care Med 196:1191–1201.

9. Pai M, Nicol MP, Boehme CC. 2016. Tuberculosis Diagnostics: State of the Art and Future Directions. Microbiol Spectr 4:TBTB2-0019-2016.

10. Coll F, Phelan J, Hill-Cawthorne GA, Nair MB, Mallard K, Ali S, Abdallah AM, Alghamdi S, Alsomali M, Ahmed AO, Portelli S, Oppong Y, Alves A, Bessa TB, Campino S, Caws M, Chatterjee A, Crampin AC, Dheda K, Furnham N, Glynn JR, Grandjean L, Minh Ha D, Hasan R, Hasan Z, Hibberd ML, Joloba M, Jones-Lopez EC, Matsumoto T, Miranda A, Moore DJ, Mocillo N, Panaiotov S, Parkhill J, Penha C, Perdigao J, Portugal I, Rchiad Z, Robledo J, Sheen P, Shesha NT, Sirgel FA, Sola C, Oliveira Sousa E, Streicher EM, Helden PV, Viveiros M, Warren RM, McNerney R, Pain A, et al. 2018. Genome-wide analysis of multi- and extensively drug-resistant Mycobacterium tuberculosis. Nat Genet 50:307–316.

11. Ng KCS, van Deun A, Meehan CJ, Torrea G, Driesen M, Gabriels S, Rigouts L, Andre E, de Jong BC. 2018. Xpert Ultra Can Unambiguously Identify Specific Rifampin Resistance-Conferring Mutations. J Clin Microbiol 56:e00686–18.

12. Tolani MP, D’Souza D T, Mistry NF. 2012. Drug resistance mutations and heteroresistance detected using the GenoType MTBDRplus assay and their implication for treatment outcomes in patients from Mumbai, India. BMC Infect Dis 12:9.

13. Canetti G, Fox W, Khomenko A, Mahler HT, Menon NK, Mitchison DA, Rist N, Smelev NA. 1969. Advances in techniques of testing mycobacterial drug sensitivity, and the use of sensitivity tests in tuberculosis control programmes. Bull World Health Organ 41:21–43.

14. Canetti G, Froman S, Grosset J, Hauduroy P, Langerova M, Mahler HT, Meissner G, Mitchison DA, Sula L. 1963. Mycobacteria: Laboratory Methods for Testing Drug Sensitivity and Resistance. Bull World Health Organ 29:565–78.

15. Blakemore R, Story E, Helb D, Kop J, Banada P, Owens MR, Chakravorty S, Jones M, Alland D. 2010. Evaluation of the analytical performance of the Xpert MTB/RIF assay. J Clin Microbiol 48:2495–501.

16. Zetola NM, Shin SS, Tumedi KA, Moeti K, Ncube R, Nicol M, Collman RG, Klausner JD, Modongo C. 2014. Mixed Mycobacterium tuberculosis complex infections and false-negative results for rifampin resistance by GeneXpert MTB/RIF are associated with poor clinical outcomes. J Clin Microbiol 52:2422–9.

17. Chakravorty S, Simmons AM, Rowneki M, Parmar H, Cao Y, Ryan J, Banada PP, Deshpande S, Shenai S, Gall A, Glass J, Krieswirth B, Schumacher SG, Nabeta P, Tukvadze N, Rodrigues C, Skrahina A, Tagliani E, Cirillo DM, Davidow A, Denkinger CM, Persing D, Kwiatkowski R, Jones M, Alland D. 2017. The New Xpert MTB/RIF Ultra: Improving Detection of Mycobacterium tuberculosis and Resistance to Rifampin in an Assay Suitable for Point-of-Care Testing. MBio 8:e00812–17.

18. Operario DJ, Koeppel AF, Turner SD, Bao Y, Pholwat S, Banu S, Foongladda S, Mpagama S, Gratz J, Ogarkov O, Zhadova S, Heysell SK, Houpt ER. 2017. Prevalence and extent of heteroresistance by next generation sequencing of multidrug-resistant tuberculosis. PLoS One 12:e0176522.

19. Eilertson B, Maruri F, Blackman A, Herrera M, Samuels DC, Sterling TR. 2014. High proportion of heteroresistance in gyrA and gyrB in fluoroquinolone-resistant Mycobacterium tuberculosis clinical isolates. Antimicrob Agents Chemother 58:3270–5.

20. Makhado NA, Matabane E, Faccin M, Pincon C, Jouet A, Boutachkourt F, Goeminne L, Gaudin C, Maphalala G, Beckert P, Niemann S, Delvenne JC, Delmee M, Razwiedani L, Nchabeleng M, Supply P, de Jong BC, Andre E. 2018. Outbreak of multidrug-resistant tuberculosis in South Africa undetected by WHO-endorsed commercial tests: an observational study. Lancet Infect Dis 18:1350–59.

21. Walker TM, Kohl TA, Omar SV, Hedge J, Del Ojo Elias C, Bradley P, Iqbal Z, Feuerriegel S, Niehaus KE, Wilson DJ, Clifton DA, Kapatai G, Ip CLC, Bowden R, Drobniewski FA, Allix-Beguec C, Gaudin C, Parkhill J, Diel R, Supply P, Crook DW, Smith EG, Walker AS, Ismail N, Niemann S, Peto TEA, Modernizing Medical Microbiology Informatics G. 2015. Whole-genome sequencing for prediction of Mycobacterium tuberculosis drug susceptibility and resistance: a retrospective cohort study. Lancet Infect Dis 15:1193–1202.

22. Vincent V, Rigouts L, Nduwamahoro E, Holmes B, Cunningham J, Guillerm M, Nathanson CM, Moussy F, De Jong B, Portaels F, Ramsay A. 2012. The TDR Tuberculosis Strain Bank: a resource for basic science, tool development and diagnostic services. Int J Tuberc Lung Dis 16:24–31.

23. Coll F, McNerney R, Guerra-Assuncao JA, Glynn JR, Perdigao J, Viveiros M, Portugal I, Pain A, Martin N, Clark TG. 2014. A robust SNP barcode for typing Mycobacterium tuberculosis complex strains. Nat Commun 5:4812.

24. Baker CN, Thornsberry C, Hawkinson RW. 1983. Inoculum standardization in antimicrobial susceptibility testing: evaluation of overnight agar cultures and the Rapid Inoculum Standardization System. J Clin Microbiol 17:450–7.

25. Donay JL, Fernandes P, Lagrange PH, Herrmann JL. 2007. Evaluation of the inoculation procedure using a 0.25 McFarland standard for the BD Phoenix automated microbiology system. J Clin Microbiol 45:4088–9.

26. Rigouts L, Gumusboga M, de Rijk WB, Nduwamahoro E, Uwizeye C, de Jong B, Van Deun A. 2013. Rifampin resistance missed in automated liquid culture system for Mycobacterium tuberculosis isolates with specific rpoB mutations. J Clin Microbiol 51:2641–5.

27. Tagliani E, Hassan MO, Waberi Y, De Filippo MR, Falzon D, Dean A, Zignol M, Supply P, Abdoulkader MA, Hassangue H, Cirillo DM. 2017. Culture and Next-generation sequencing-based drug susceptibility testing unveil high levels of drug-resistant-TB in Djibouti: results from the first national survey. Sci Rep 7:17672.

28. El Achkar S, Demanche C, Osman M, Rafei R, Ismail MB, Yaacoub H, Pincon C, Duthoy S, De Matos F, Gaudin C, Trovato A, Cirillo DM, Hamze M, Supply P. 2019. Drug-Resistant Tuberculosis, Lebanon, 2016 - 2017. Emerg Infect Dis 25:564–568.

29. Kohl TA, Utpatel C, Schleusener V, De Filippo MR, Beckert P, Cirillo DM, Niemann S. 2018. MTBseq: a comprehensive pipeline for whole genome sequence analysis of Mycobacterium tuberculosis complex isolates. PeerJ 6:e5895.

30. Lawn SD, Nicol MP. 2011. Xpert(R) MTB/RIF assay: development, evaluation and implementation of a new rapid molecular diagnostic for tuberculosis and rifampicin resistance. Future Microbiol 6:1067–82.

31. Hain Lifescience. 2015. GenoType MTBDR*plus* VER 2.0 IFU-304A-09 Molecular Genetic Assay for Identification of the *M. tuberculosis* Complex and its Resistance to Rifampicin and Isoniazid from Clinical Specimens and Cultivated Samples.1–14.

32. Andre E, Isaacs C, Affolabi D, Alagna R, Brockmann D, de Jong BC, Cambau E, Churchyard G, Cohen T, Delmee M, Delvenne JC, Farhat M, Habib A, Holme P, Keshavjee S, Khan A, Lightfoot P, Moore D, Moreno Y, Mundade Y, Pai M, Patel S, Nyaruhirira AU, Rocha LE, Takle J, Trebucq A, Creswell J, Boehme C. 2016. Connectivity of diagnostic technologies: improving surveillance and accelerating tuberculosis elimination. Int J Tuberc Lung Dis 20:999–1003.

33. Metcalfe JZ, Streicher E, Theron G, Colman RE, Penaloza R, Allender C, Lemmer D, Warren RM, Engelthaler DM. 2017. Mycobacterium tuberculosis Subculture Results in Loss of Potentially Clinically Relevant Heteroresistance. Antimicrob Agents Chemother 61.

34. Horne DJ, Kohli M, Zifodya JS, Schiller I, Dendukuri N, Tollefson D, Schumacher SG, Ochodo EA, Pai M, Steingart KR. 2019. Xpert MTB/RIF and Xpert MTB/RIF Ultra for pulmonary tuberculosis and rifampicin resistance in adults. Cochrane Database Syst Rev 6:CD009593.

35. Luetkemeyer AF, Kendall MA, Wu X, Lourenco MC, Jentsch U, Swindells S, Qasba SS, Sanchez J, Havlir DV, Grinsztejn B, Sanne IM, Firnhaber C, Adult ACTGAST. 2014. Evaluation of two line probe assays for rapid detection of Mycobacterium tuberculosis, tuberculosis (TB) drug resistance, and non-TB Mycobacteria in HIV-infected individuals with suspected TB. J Clin Microbiol 52:1052–9.

36. Pai M, Behr MA, Dowdy D, Dheda K, Divangahi M, Boehme CC, Ginsberg A, Swaminathan S, Spigelman M, Getahun H, Menzies D, Raviglione M. 2016. Tuberculosis. Nat Rev Dis Primers 2:16076.

37. Rigouts L, Coeck N, Gumusboga M, de Rijk WB, Aung KJ, Hossain MA, Fissette K, Rieder HL, Meehan CJ, de Jong BC, Van Deun A. 2016. Specific gyrA gene mutations predict poor treatment outcome in MDR-TB. J Antimicrob Chemother 71:314–23.

